# Smartphone-Validated Portable Paper-Based Device Integrated with an Electropolymerized Molecularly Imprinted Polymer for Serotonin Detection in Serum Samples

**DOI:** 10.64898/2026.06.30.735496

**Authors:** Harshit Borasi, Bhagyesh Parmar, Pranjal Agarwal, Dhiraj Bhatia, Amit K. Yadav

## Abstract

Accurate and decentralized quantification of serotonin, also known as 5-hydroxytryptamine (5-HT), in biological fluids is critically important for the diagnosis, prognosis, and therapeutic monitoring of neurological and psychiatric disorders. However, conventional analytical methods generally rely on centralized laboratory infrastructure, skilled personnel, and labor-intensive sample processing, which restrict their applicability in rapid near-patient and point-of-care settings. Herein, we report a portable molecularly imprinted polymer (MIP)-based electrochemical sensing platform for selective and on-site detection of serotonin using screen-printed carbon electrodes (SPCEs). The biomimetic recognition interface was fabricated through direct electropolymerization of a polydopamine recognition layer in the presence of serotonin as the template molecule, followed by template extraction to generate complementary recognition cavities for selective rebinding. The sensor fabrication parameters, including monomer concentration, electropolymerization cycles, template-to-monomer stoichiometry, and electrolyte pH, were systematically optimized to achieve improved sensitivity, selectivity, and signal stability. Under optimized conditions, the MIP/SPCE sensor exhibited a broad linear response from 10 pM −10 µM in phosphate buffer, with a correlation coefficient of R^2^ = 0.974 and an ultralow limit of detection of 0.16 pM. The analytical applicability of the platform was further validated in spiked artificial serum, where the sensor achieved an LOD of 0.12 pM, satisfactory recovery values of 88.66-96.02%, and acceptable precision with RSD values ≤ 8.43% (n=3), confirming its reliability in a complex biological matrix. The developed sensor demonstrated excellent selectivity toward serotonin against physiologically relevant interferents, maintaining signal retention between 99% and 101%. In addition, the platform showed high operational repeatability with an RSD of 0.45%, good inter-electrode reproducibility with an RSD of 6.3%, and long-term storage stability, retaining 90-110% of its initial response over 28 days. Importantly, cross-platform validation using a smartphone-coupled potentiostat demonstrated strong analytical agreement with laboratory-grade instrumentation, as evidenced by R^2^ = 0.9967 and a slope of 1.023. These findings establish the proposed MIP/SPCE platform as a simple, low-cost, portable, and smartphone-compatible electrochemical device for field-deployable serotonin monitoring in clinically relevant samples.

## 1. Introduction

Serotonin, also known as 5-hydroxytryptamine (5-HT), is a biogenic monoamine neurotransmitter primarily synthesized in presynaptic neurons of the Central Nervous System (CNS) [1]. However, more than 90% of the body’s total Serotonin is stored in platelets, which serve as primary storage sites by readily taking up the amine from the plasma [2,3]. It serves as a critical regulatory molecule across multiple physiological systems, involved in the regulation of mood, behavior, memory, and other physiological processes, including gastro-intestinal homeostasis, bladder control, and cardiovascular function [4,5]. Dysregulation of serotonergic homeostasis is directly implicated in the pathophysiology of major depressive disorder, anxiety disorders, autism spectrum disorder, and Alzheimer’s [6,7]. Serotonin syndrome results from excessive serotonergic activity due to the use of therapeutic medicine, abuse of recreational drugs, or intentional overdoses, with symptoms ranging from altered mental status, autonomic dysfunction, and neuromuscular hyperactivity [8,9]. Thus, accurate, timely quantification of Serotonin is indispensable for early diagnosis, therapeutic drug monitoring, and mechanistic neurochemical studies. Conventional serotonin assays rely predominantly on high-performance liquid chromatography (HPLC) [10], liquid chromatography (LC) combined with mass spectrometry [11,12], ultra-high-performance liquid chromatography (UHPLC) with fluorescence detection (FD) [13], or tandem mass spectrometry (UHPLC-MS/MS) [14–16], high-precision liquid chromatography with an electrochemical detector (HPLC-ECD) [17], gas chromatography coupled with Mass Spectrometry (GC-MS) [18], spectrophotometry [19–22] and capillary electrophoresis [23]. While these platforms deliver excellent analytical figures of merit, they are fundamentally incompatible with point-of-care deployment. They require centralized laboratory infrastructure, trained personnel, and sophisticated sample preparation protocols.

Electrochemical biosensors offer a compelling alternative, combining rapid response, high sensitivity, wide linear range, cost-effectiveness, accuracy, ease of operation and manufacturing, and their point-of-care employability [24,25]. Various electrochemical sensing platforms have been developed for the quantitative determination of Serotonin, leveraging its inherent electroactivity via voltammetric techniques such as cyclic voltammetry (CV) and differential pulse voltammetry (DPV) [26]. These systems have been implemented across a range of electrodes, encompassing carbon paste electrodes (CPE) [27–31], glassy carbon electrodes (GCE) [32–34], screen-printed carbon electrodes (SPCE) [35,36], and graphite pencil electrodes [37,38], each offering distinct physicochemical and fabrication advantages relevant to analytical performance. Various factors limit the sensitivity and accuracy of electrochemical sensing of Serotonin. The presence of electroactive biomolecules in biological samples, such as ascorbic acid (AA) and uric acid (UA), which are present at higher concentrations than Serotonin and have similar oxidation potentials, compromises the selectivity of the sensing platform [39,40]. Second, Serotonin and structurally similar phenolic amines undergo oxidation and strong adsorption on carbon surfaces, leading to electrode fouling and signal drift under repeated use [26]. Third, due to the high adsorption rate of Serotonin by platelets, levels of Serotonin in human samples become too low to detect, resulting in poor sensitivity and selectivity [5]. Modification of electrodes with conductive or catalytic materials can circumvent these challenges, improving the sensitivity and selectivity of serotonin detection. Modifying the electrodes with conducting polymers [41], carbon nanomaterials [42,43], metal nanoparticles [44,45], and ionic liquids [46] has been demonstrated to improve serotonin detection. Molecularly imprinted polymers (MIPs) offer a promising approach to improve the selectivity and sensitivity of analyte detection [47,48]. Molecularly imprinted polymers (MIPs) provide a powerful strategy to introduce molecular recognition into fully synthetic materials by creating template-specific cavities complementary in shape and functional group arrangement to the target analyte [49–51].

In this work, we developed a polydopamine (PDP) MIP electrochemical sensor for highly sensitive, rapid, and selective detection of Serotonin. The presence of catechol and aminoethyl side chains in dopamine provides essential functional groups for establishing non-covalent interactions with template molecules. These groups facilitate the formation of hydrogen bonds and Ö-Ö stacking, which are critical for stabilizing the prepolymerization complex and ultimately yielding a molecularly imprinted polymer (MIP) with high selectivity [52]. Polydopamine has been widely employed for the detection of small molecules [52–54], proteins [55,56], bacteria [57,58], and viruses [59,60] due to its ability to form well-defined, stable cavities. Moreover, dopamine can be polymerized in a potential window that does not overlap with the oxidation potential window of Serotonin [53,54]. Screen-printed electrodes (SPCEs) were used to fabricate our PDP-MIP/SPCE platform. The cost-effectiveness and suitability for large-scale manufacturing, combined with their disposable nature and customizable surface chemistry, make SPCEs ideal candidates for analytical applications [55]. These platforms offer significant advantages for point-of-care applications for their disposability and amenability to miniaturization and integration into portable devices. Despite substantial progress in electrochemical serotonin sensing, the simultaneous achievement of sub-picomolar detection, robust selectivity in complex biological matrices, and portability has not been demonstrated on a single integrated platform [56–60]. In this work, we address these gaps by developing a PDP-based molecularly imprinted electrochemical sensor on SPCEs and systematically evaluating its performance from controlled buffer conditions through to spiked artificial serum, with the entire analytical workflow validated on PalmSens Sensit Smart, a smartphone-operated portable potentiostat. To the best of our knowledge, this is the first report of a polydopamine-based molecularly imprinted sensor on a screen-printed carbon electrode achieving sub-picomolar serotonin detection in a spiked biological matrix, with quantitative cross-platform validation confirming analytical equivalence between benchtop and smartphone-operated instrumentation on a single disposable sensing interface.

## 2. Experimental Section

### 2.1 Materials and Methods

Dopamine hydrochloride was purchased from Alfa Aesar. Serotonin hydrochloride and histamine were obtained from Sigma-Aldrich. Tris base was procured from BIOSCIENCE, and L-tryptophan was purchased from HiMedia. Anhydrous D-glucose, potassium chloride, anhydrous sodium phosphate monobasic, sodium phosphate dibasic dihydrate, urea, sodium chloride, potassium ferricyanide, potassium ferrocyanide, glycine, and adenine were all sourced from Sisco Research Laboratories (SRL, India). Screen-printed carbon electrodes (Design ID: p1067-GA-0011_vA) were purchased from Fle Medical Solutions. All reagents were of analytical grade and utilized as received without subsequent purification. Aqueous solutions were universally formulated using Milli-Q water, and all experimental procedures were conducted at ambient room temperature (25 °C). A supporting electrolyte comprising 0.2 M PBS and 5.0 mM [Fe(CN)_6_]^3−/4−^ was prepared at pH (6.0, 6.5, 7.0, 7.5, and 8.0). To systematically evaluate the electrochemical sensitivity of the fabricated MIP sensor, a 1.0 mM serotonin stock solution was diluted in a series to generate working standards spanning 1 pM to 100 µM.

### 2.2 Characterization Techniques

To confirm the sequential modification steps, the surface chemistries of the non-imprinted polymer (PDP-NIP), the intermediate PDP/serotonin complex, and the target-extracted molecularly imprinted polymer (PDP-MIP) were elucidated via Fourier transform infrared spectroscopy (FTIR) using Bruker Invenio-S FTIR. Additionally, the surface topographies of the PDP-NIP and PDP-MIP films were evaluated using a Bruker Nano Wizard Sense AFM. Electrochemical measurements for optimization and full analytical characterization were performed on a PalmSens EmStat4S and a PalmSens Sensit Smart smartphone-operated potentiostat, using PSTrace and PSTouch software, respectively. A supporting electrolyte of 0.2 M phosphate-buffered saline (PBS) containing 5.0 mM [Fe(CN)_6_]^3−/4−^ was employed for all electrochemical measurements, prepared at pH 7.0 unless otherwise stated.

### 2.3 Polydopamine MIP and NIP synthesis

Prior to surface functionalization, bare SPCEs were cleaned sequentially with ethanol and deionized water, and then electrochemically activated in 0.1 M H_2_SO_4_ by cyclic voltammetry (CV) over 10 consecutive cycles from −0.8 V to +0.8 V until a stable, reproducible cyclic voltammogram was obtained. To construct the molecularly imprinted interface, a prepolymerization solution containing 2.1 mM dopamine and 0.35 mM serotonin (1:6 template-to-monomer molar ratio) was prepared in 0.1 M phosphate buffer (PB, pH 8.0). The solution was incubated in the dark for 1 hour to allow template-monomer pre-assembly. A 50 μL aliquot was cast onto the activated electrode surface, and electropolymerization was induced by CV over a potential window of −0.5 V to +0.5 V for 10 cycles at 50 mV/s [53]. A non-imprinted polymer control (PDP-NIP) was deposited under identical conditions, omitting Serotonin from the prepolymerization mixture. All functionalized electrodes were rinsed with deionized water following polymerization.

Template extraction was performed by continuous electrochemical cycling in 0.1 M PB (pH 8.0) between −0.2 V and +0.85 V for 10 cycles at 50 mV/s, which disrupted the non-covalent template-polymer interactions and liberated Serotonin from the imprinted cavities without compromising the polymeric framework [53]. To maximize analytical performance, key fabrication variables were systematically optimized using differential pulse voltammetry (DPV) in the presence of the [Fe(CN)_6_]^3−/4−^ redox probe. The parameters evaluated were: (i) dopamine monomer concentration (1.0-5.0 mM), (ii) number of electropolymerization cycles (5-30), (iii) template-to-monomer molar ratio (1:3, 1:6, 1:9, and 1:12), and (iv) electrolyte pH (6.0-8.0). Each variable was independently optimized while keeping all others at their established optimal values (Fig.1)

**Figure 1:**
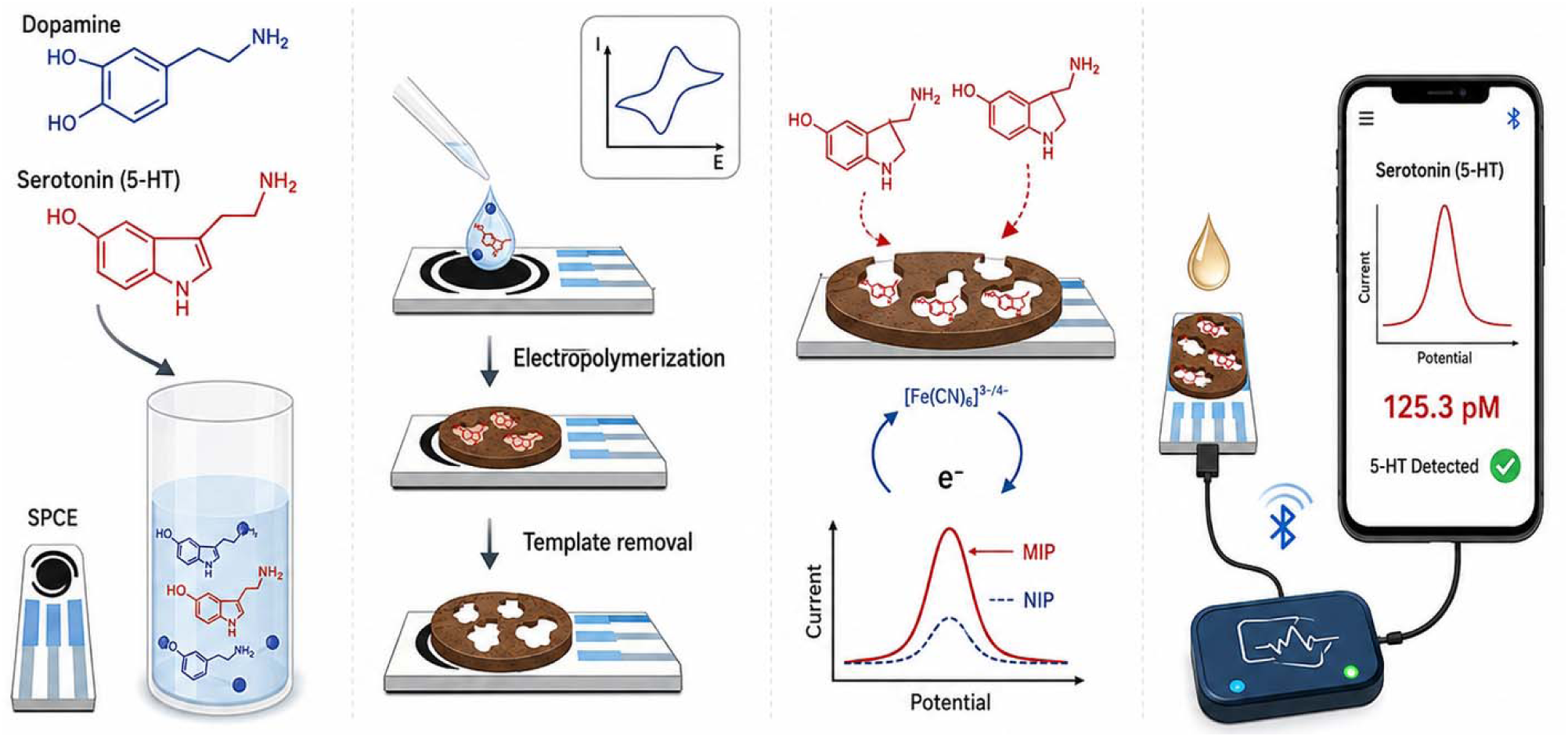
Schematic illustration of the fabrication and portable electrochemical detection strategy of the molecularly imprinted SPCE sensor for serotonin (5-HT).

### 2.4 Spiked Sample Preparation

To validate analytical performance in a complex biological matrix, serotonin detection was evaluated in artificial human serum. Two distinct sample sets were prepared: undiluted serum and a 1:10 serum dilution, each spiked with serotonin standard solutions to yield a concentration gradient spanning 1 pM to 100 μM. For electrochemical measurements, 10 μL aliquots of the spiked serum preparations were added to 50 μL of [Fe(CN)_6_]^3−/4−^ working buffer in 0.1 M PBS

## 3. Results and Discussion

### 3.1 FT-IR spectroscopy

FTIR spectroscopy was employed to confirm precursor signatures, polymer formation, template incorporation, and subsequent removal. The spectra of dopamine and Serotonin exhibited distinct features reflecting their structural differences. Dopamine showed prominent absorption in the 3200-3300 cm^−1^ region corresponding to O-H stretching of catechol groups, along with bands in the 1240-1250 cm^−1^ region attributed to C-O stretching (Figure 2.i.) [61–63]. In contrast, Serotonin displayed a characteristic band at ∼1526 cm^−1^, assigned to indole-associated vibrations, which is absent in dopamine and serves as a diagnostic marker for Serotonin [64]. All polymeric samples (NIP, MIP, and MIP Ser) exhibited a broad band in the 3400-3200 cm^−1^ region (O-H/N-H stretching), along with characteristic peaks at ∼1680-1600 cm^−1^ (C=O/C=N stretching) and ∼1240-1090 cm^−1^ (C-O and C-O-C vibrations), confirming the formation of a polydopamine network via oxidative polymerization of dopamine. A distinct peak at ∼1526 cm^−1^, corresponding to Serotonin, was clearly observed in the MIP Ser, confirming successful incorporation of the template during polymerization. This peak was absent in the NIP, indicating that the polymer backbone originates solely from dopamine. Following template removal, the MIP spectrum showed complete disappearance of the serotonin-specific bands (∼1526 cm^−1^ and ∼1380 cm^−1^).

**Figure 2:**
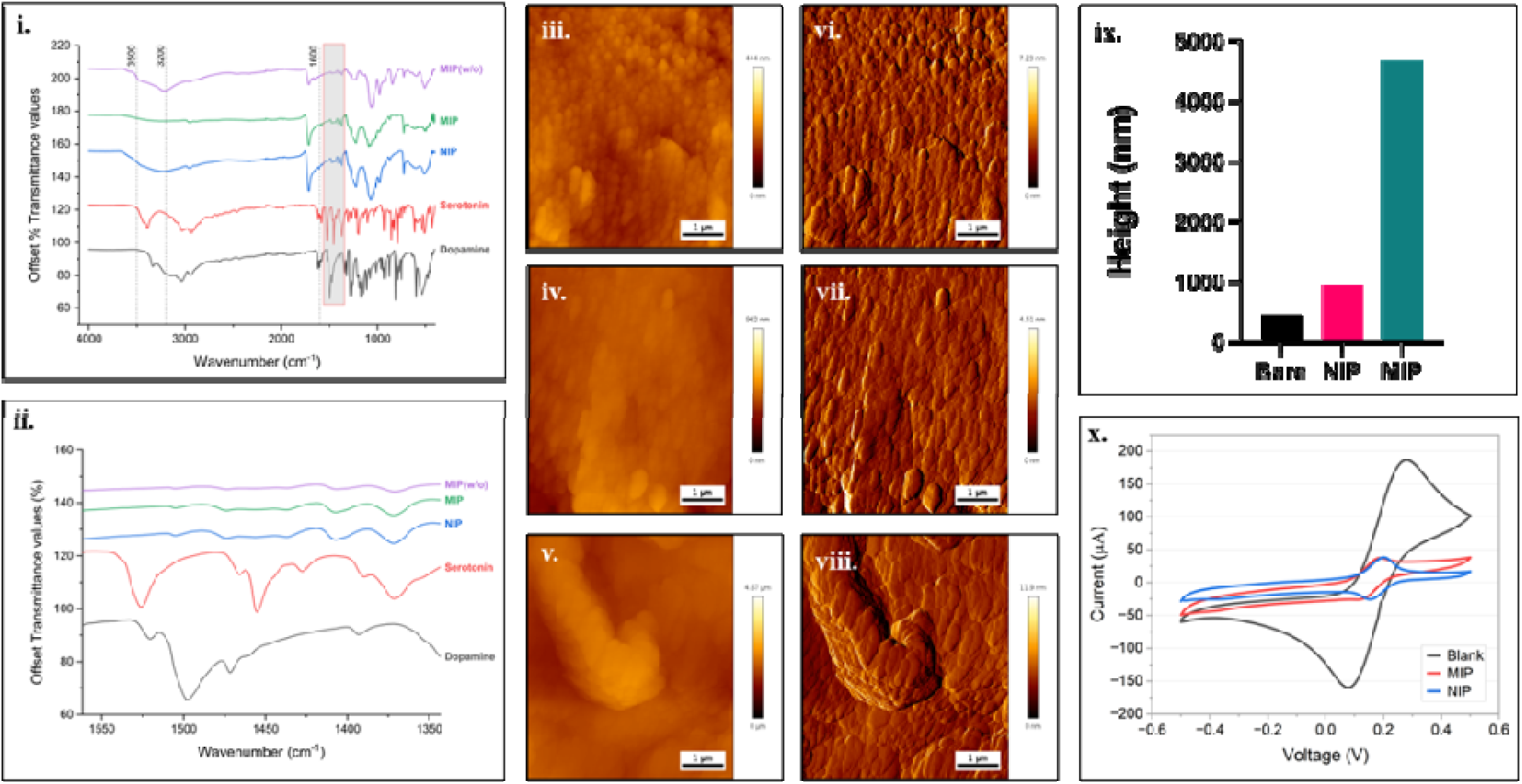
Spectroscopic, morphological, and electrochemical characterization of the PDP-MIP-modified SPCE surface. *(i)* Fourier-transform infrared (FTIR) spectra of dopamine, Serotonin, non-imprinted polymer (NIP), molecularly imprinted polymer (MIP), and MIP prior to template removal (MIP w/o). *(ii)* Magnified FTIR spectra (1350-1550 cm^−1^) highlighting variations in C-N stretching and aromatic ring vibrations, where subtle shifts in peak positions and intensities indicate intermolecular interactions during polymer formation and successful imprinting. *(iii-v)* Atomic force microscopy (AFM) 2D topographical images of *(iii)* bare SPCE, *(iv)* NIP-modified electrode, and *(v)* MIP-modified electrode (after template removal). *(vi-viii)* Corresponding 3D AFM topography images of *(vi)* bare SPCE, *(vii)* NIP, and *(viii)* MIP surfaces, where the MIP demonstrates enhanced surface roughness and pronounced topographical features compared to the NIP, reflecting the formation of porous structures and accessible binding sites. *(ix)* Quantita ive surface roughness/height analysis derived from AFM data, showing increased surface height variation for the MIP compared to the NIP and bare electrode, further confirming the formation of a rough and porous imprinted surface. *(x)* Cyclic voltammetry (CV) responses of bare SPCE (blank), NIP-, and MIP-modified electrodes.

### 3.2 AFM study

Surface topography and phase imaging (Fig. 2. iii-viii) confirm the successful transition from the granular bare SPCE to a continuous, nanoparticulate polydopamine (PDP) architecture examined using contact mode and a 1 µm × 1 µm scan area. The PDP-MIP surface (Fig. 2.v.) exhibits higher topographical complexity and porosity compared to the smoother non-imprinted (NIP) control (Fig. 2.iv.), indicating the formation of accessible imprinted cavities. Quantitative height analysis reveals a significant increase in surface elevation following modification (Fig. 2.ix.). While the bare SPCE features a relatively low profile, the MIP-functionalized surface shows substantial vertical growth (reaching approximately 4 µm in the primary scanning area), reflecting the deposition of a robust polymeric matrix.

### 3.3 Electrochemical Studies

#### 3.3.1 pH Study

The electrochemical dynamics of the MIP sensor and the specific binding affinity between the target analyte and the molecularly imprinted polymer (MIP) are fundamentally governed by the pH of the supporting electrolyte [52]. Consequently, the electroanalytical performance of the MIP-modified screen-printed carbon electrode (SPCE) was systematically evaluated in phosphate-buffered saline (PBS) containing a [Fe(CN)_6_]^3−/4−^ redox probe across a pH gradient of 6.0 to 8.0, in the potential range of −0.5 to 0.5V. Differential pulse voltammetry (DPV) profiles (Fig. 3.iv.) showed a progressive increase in peak current as the electrolyte pH was raised from 6.0 to 7.0, followed by a distinct decrease in peak current at pH 8.0. This suggests that the specific non-covalent interactions between Serotonin and the complementary microenvironments within the polydopamine recognition cavities are thermodynamically optimized at pH 7.0. Because the PDP-MIP/SPCE platform exhibited its highest electrocatalytic capacity at pH 7.0, PBS formulated at this pH with the [Fe(CN)_6_]^3−/4−^ redox system was adopted as the standardized medium for all subsequent quantitative evaluations.

**Fig. 3.**
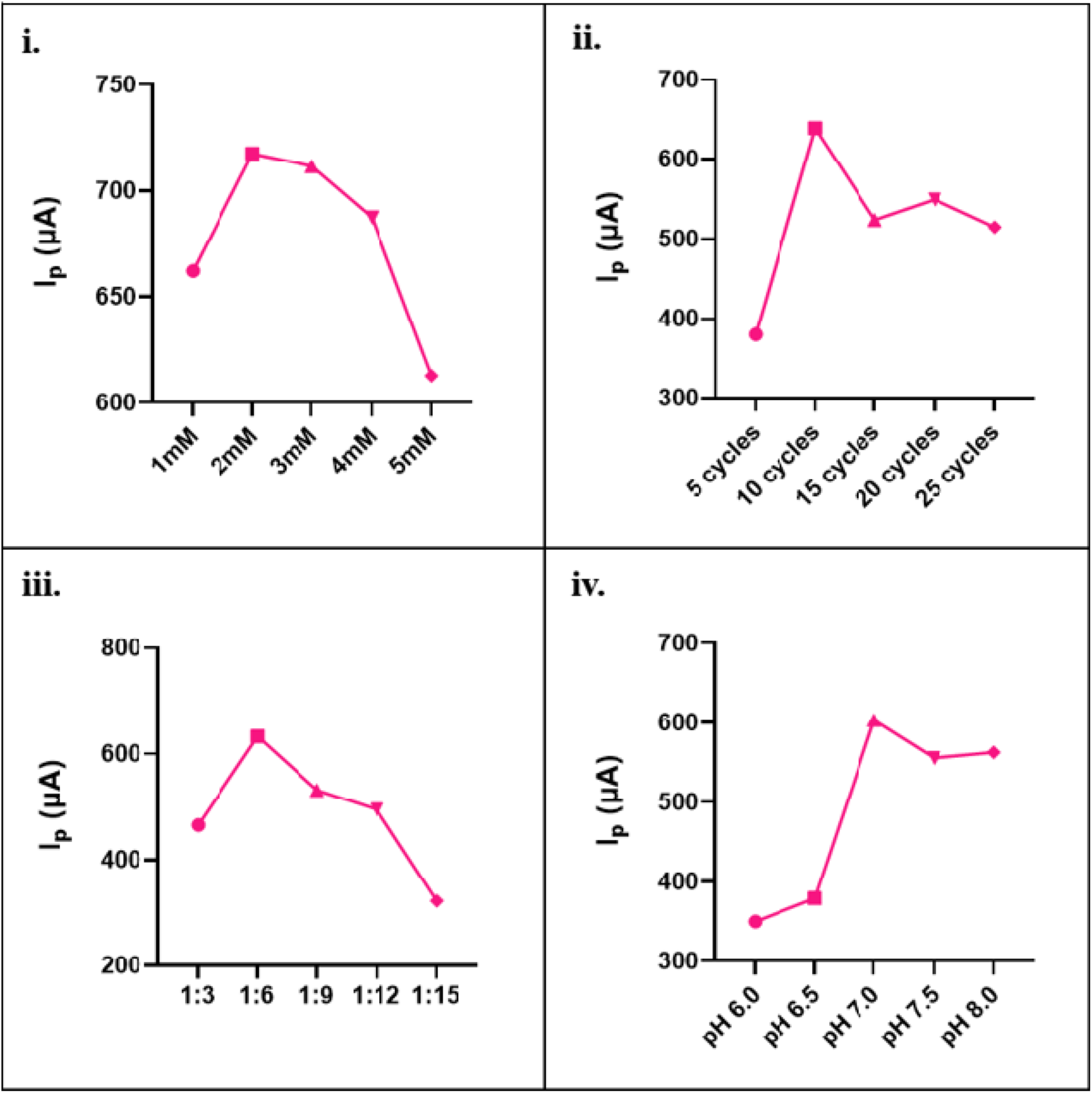
Optimization of experimental parameters for the PDP-MIP/SPCE sensor performance. Optimization of experimental parameters for PDP-MIP/SPCE sensor performance. (i). Effect of dopamine monomer concentration (1.0-5.0 mM) on DPV peak current response, showing an optimum at 2.0 mM. (ii). Influence of the number of electropolymerization cycles on sensor response, demonstrating a maximum at 10 cycles, followed by signal attenuation due to mass transport limitations at higher cycle numbers. (iii) Effect of template-to-monomer molar ratio on electrochemical response, with optimal performance at 1:6 and declining signals at higher monomer excess due to matrix over-crosslinking. (iv) Effect of supporting electrolyte pH (6.0–-.0) on DPV peak current, with peak sensitivity achieved at pH 7.0

#### 3.3.2 Optimal Fabrication Conditions

To maximize the analytical performance of the PDP-MIP/SPCE, the parameters governing its electrosynthesis and operation were systematically optimized (Fig. 3). The dopamine monomer concentration was varied from 1.0 to 5.0 mM, with the Faradaic peak current of the [Fe(CN)_6_]^3−/4−^redox probe reaching a maximum at 2.0 mM (Fig. 3.i.) [65]. Beyond this concentration, the current response declined, most likely due to film aggregation and restricted diffusional pathways resulting from an overly dense polymer network. A concentration of 2.1 mM was accordingly adopted for all subsequent fabrications [53].

The effect of the electropolymerization cycle number was evaluated from 5 to 30 cycles using 2.1 mM dopamine in 0.1 M PB (pH 8.0) [52]. The maximum DPV current density was achieved at 10 cycles (Fig. 3.ii). Beyond this point, the progressive thickening of the polymeric film imposed mass-transport limitations and partially buried the imprinted cavities, diminishing both charge-transfer efficiency and recognition-te accessibility [53]. Similarly, the template-to-monomer stoichiometric ratio was screened at 1:3, 1:6, 1:9, and 1:12. The peak serotonin oxidation signal was observed at a 1:6 ratio (Fig. 3.iii.); higher monomer proportions 1:9 and 1:12) yielded diminished responses, attributable to an over-cross-linked, less conductive matrix that impedes target diffusion and reduces the ratio of occupied to total cavities [52,53].

#### 3.3.3 Scan Rate Study

Scan rate-dependent CV measurements were performed from 0.005 to 0.15 V/s to investigate the electrochemical kinetics of PDP-MIP/SPCE in 0.2 M PBS containing 5 mM [Fe(CN)O]^3−/4−^ in a potential window of −0.5 V to +0.5 V (Fig. 4.i.). The PDP-MIP-modified electrode exhibited a well-defined, reversible redox couple with anodic and cathodic peaks at +0.210 V and +0.140 V (vs. Ag/AgCl), respectively, confirming that the PDP layer supports efficient interfacial electron transfer while permitting redox probe permeation (Fig. 4.i.). The unmodified SPCE showed no characteristic Faradaic features within the selected potential window (Fig. 4.ii.), and the PDP-NIP electrode exhibited an analogous redox couple to the MIP, as expected from equivalent polymer backbone chemistry. Both anodic and cathodic peak currents scaled linearly with scan rate (R^2^> 0.99) (Fig. 4.iii.), and a double-logarithmic analysis of peak current versus scan rate yielded slopes of 0.69 and 0.72 for the anodic and cathodic processes, respectively (Fig. 4.iv.). The slope (∼0.7) indicates adsorption-diffusion mixed control, which is characteristic of polymer-modified electrodes where mass transport through the imprinted matrix and specific binding at surface sites jointly contribute to the electrochemical response [66,67]. This conclusion is further reinforced by the higher correlation coefficient obtained for the I□ vs. ν plot (R^2^ ≈ 0.9997) compared to the I□ vs. ν^1/2^ plot (R^2^≈ 0.996). The near-constant peak-to-peak separation (ΔE□) observed across the scan rate range is characteristic of a kinetically facile, reversible electron transfer process at the polymer-electrode interface.

**Fig. 4.**
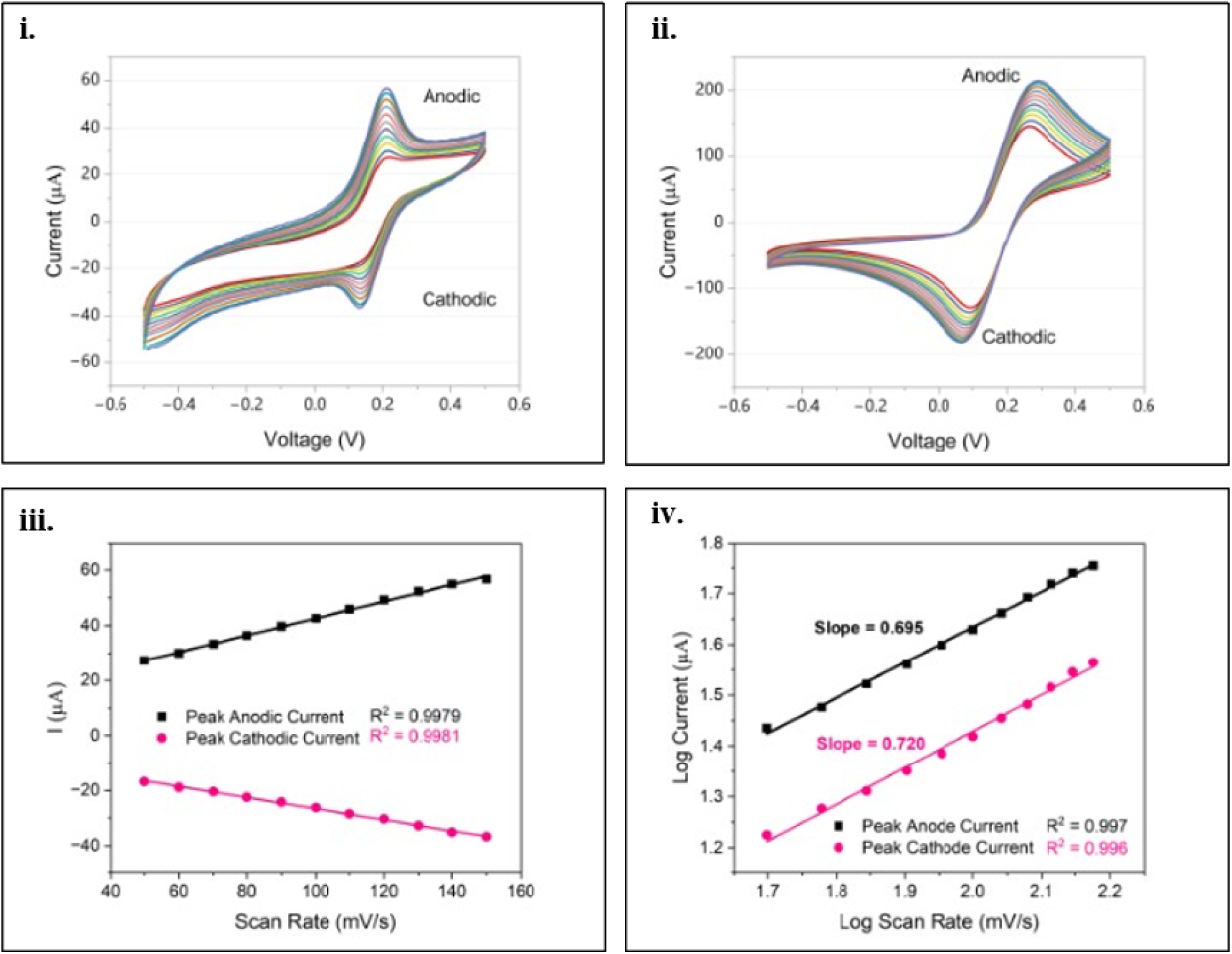
Scan rate-dependent electrochemical behavior of the PDP-MIP/SPCE sensor. *(i)* Cyclic voltammetry (CV) responses of the MIP-modified electrode recorded at increasing scan rates, showing a systematic increase in both anodic and cathodic peak currents. *(ii)* CV responses of the blank electrode under identical conditions, exhibiting lower current response and absence of specific electrochemical features compared to the MIP. *(iii)* Linear relationship between peak current (anodic and cathodic) and scan rate for the MIP sensor, indicating a surface-controlled electrochemical process. *(iv)* Log–log plot of peak current versus scan rate for the MIP electrode.

Furthermore, the electroactive surface coverage (Γ) of the adsorbed ionic species was quantified via the Brown-Anson model:

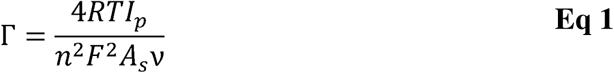

where *R* is the universal gas constant (8.314 J mol□¹ K□¹), *T* is the absolute temperature (300 K), and *F* is the Faraday constant (96,485 C mol□¹). The calculated surface coverages were 3.54 × 10□□ mol cm^−2^ for the PDP-MIP/SPCE and 3.21 × 10□□ mol cm^−2^ for the control PDP-NIP/SPCE. The elevated coverage on the MIP interface further substantiates the successful formation of geometrically and chemically complementary cavities upon serotonin extraction. These cavities inherently amplify the accessible surface area, thereby augmenting the overall electrocatalytic capacity of the imprinted matrix relative to the non-imprinted control.

The mass transport kinetics at the electrode-electrolyte interface were evaluated by determining the apparent diffusion coefficient (*D*) of the electroactive species. This was quantified using the Randles-Sevcik equation:

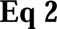

where *I_p_* denotes the faradaic peak current (anodic or cathodic), *n* represents the number of electrons transferred per redox event (*n*=2), *As* is the electroactive surface area of the SPCE (0.126 cm²), *C* is the bulk concentration of the redox probe, and ν is the applied voltammetric scan rate (V s□¹). The apparent diffusion coefficients were calculated to be 2.05 × 10□□ cm² s□¹ for the PDP-MIP/SPCE and 1.68 × 10□□ cm² s□¹ for the PDP-NIP/SPCE. The enhanced *D* value for the imprinted sensor indicates a more rapid influx of the redox probe to the electrode surface, a direct consequence of the highly porous architecture generated by the elution of the serotonin template [68].

#### 3.3.4 Electrochemical Response Study

The analytical performance of the PDP-MIP/SPCE for serotonin quantification was evaluated utilizing DPV in a 0.1 M PBS containing [Fe(CN)_6_]^3−/4−^ over a potential window of −0.5 V to 0.5 V. Standard serotonin solutions ranging from 1 pM to 100 μM were introduced as 10 μL aliquots onto the sensor surface, followed by a 10-second incubation to allow target-cavity complexation prior to voltammetric readout.

A monotonic attenuation of the DPV peak current was observed with increasing serotonin concentration (Fig. 5.i.). This signal suppression is mechanistically attributable to the progressive occupation of imprinted cavities by Serotonin. Upon rebinding, cooperative non-covalent interactions—principally hydrogen bonding between the serotonin hydroxyl/amino groups and the PDP catechol/amine functionalities, and π–π stacking between the serotonin indole ring and the aromatic regions of the PDP matrix—physically obstruct the diffusion pathways for [Fe(CN)□]³□/□□ to the underlying SPCE surface, reducing the measured Faradaic current in proportion to the number of occupied sites.

**Fig. 5.**
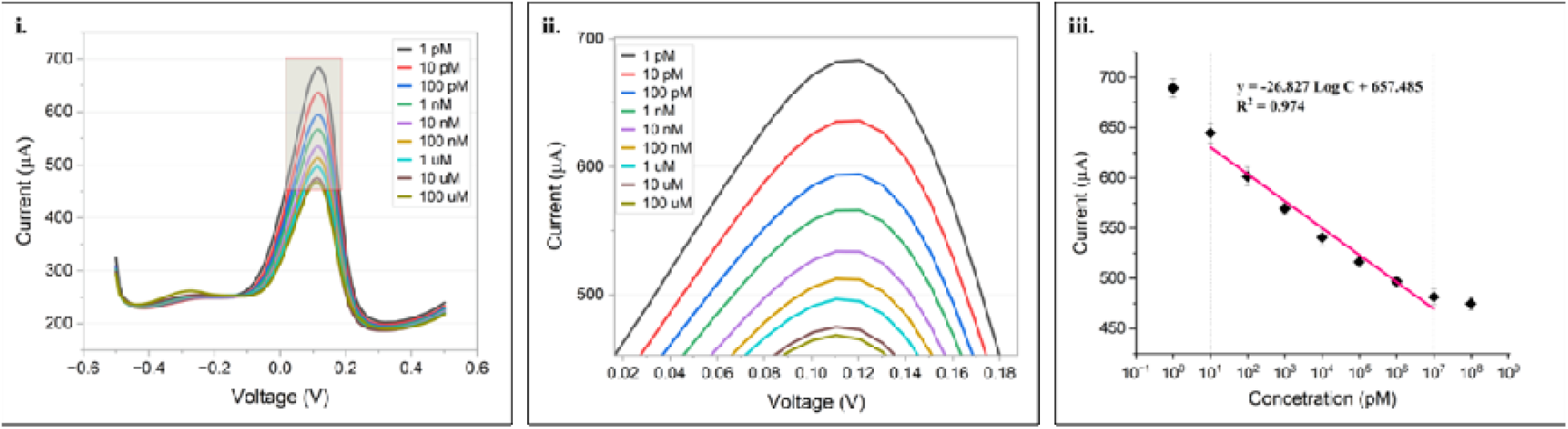
Determination of the limit of detection (LOD) of the PDP-MIP/SPCE sensor for Serotonin. *(i)* Differential pulse voltammetry (DPV) responses of the MIP-modified electrode toward increasing concentrations of Serotonin (1 pM–100 µM), showing a systematic decrease in peak current with increasing analyte concentration. *(ii)* Enlarged view of the DPV responses highlighting distinct and well-resolved peak variations at low concentration levels. *(iii)* Calibration plot of peak current versus logarithmic concentration of Serotonin, exhibiting a linear relationship within the defined range, with the regression equation. Error bars represent standard deviation (n = 3).

A robust linear correlation between peak current and logarithm of serotonin concentration was established over the range 10 pM to 10 μM (R² = 0.974), described by the regression equation:

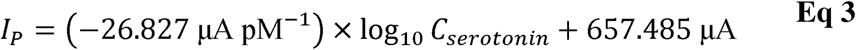

The normalized analytical sensitivity, derived by dividing the slope by the geometric electrode area (0.126 cm²), was 212.9 μA (log pM)□¹ cm□². Furthermore, the limit of detection (LOD) was calculated using the formula given in equation (2) [69], where σ represents the standard deviation of the blank electrode response, x denotes the lowest concentration measured, and s_semitog_ denotes the slope of the semilog graph.

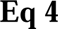

Based on these parameters, the LOD was determined to be 0.16 pM. Collectively, these metrics validate the efficacy of the engineered PDP-MIP/SPCE, characterized by a rapid temporal response, a broad linear dynamic range (10–10^7^ pM), trace-level detectability, and high sensitivity. Subsequently, the Limit of Quantification (LOQ) is found out to be 5.33 pM. The PDP-MIP/SPCE platform thus achieves a detection limit approximately two to three orders of magnitude lower than most comparable electrochemical approaches. The comparison with recent literature is summarized in Table 1.

**Table 1.**
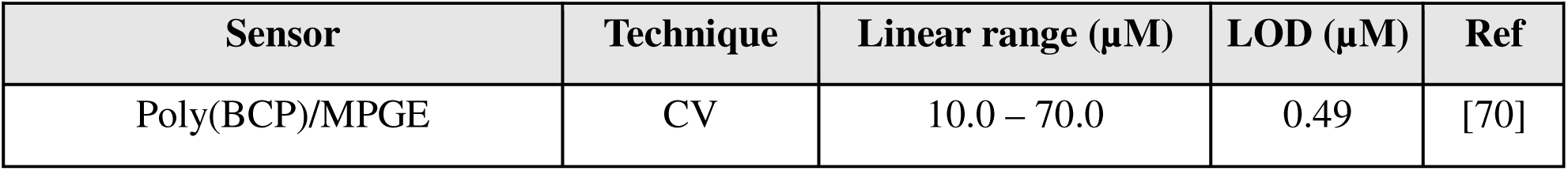
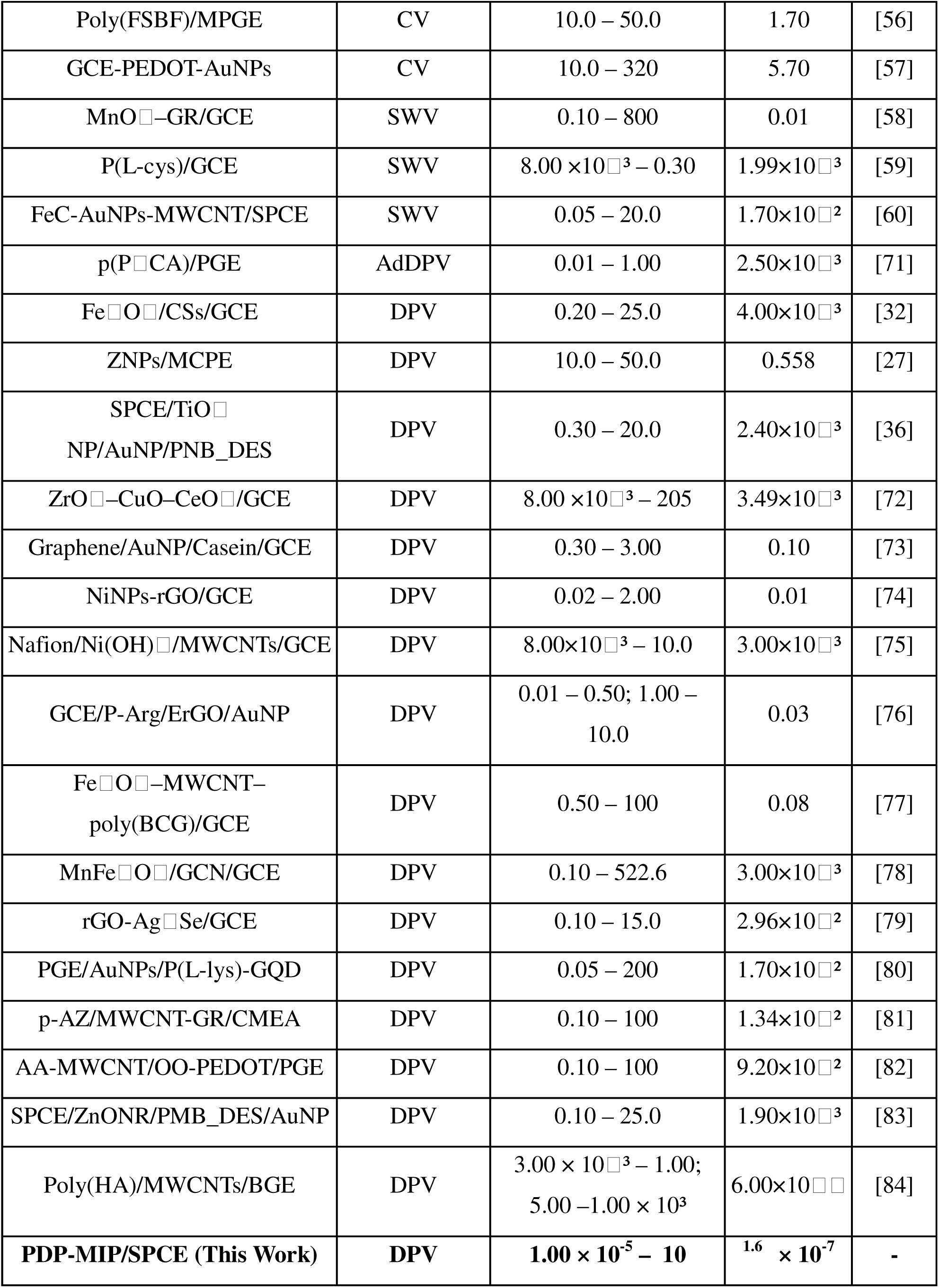
Comparison of the analytical performance of the proposed PDP/SPCE sensor with other previously reported electrochemical sensors for serotonin detection.

#### 3.3.5 Control Study

To validate the imprinting effect and ascertain the specific recognition capabilities of the proposed platform, control experiments were executed using the non-imprinted PDP-NIP/SPCE interface. The electrochemical response of the control sensor was investigated across an identical concentration gradient of Serotonin. As illustrated in Figure 6, the incremental addition of Serotonin elicited negligible fluctuations in the voltammetric peak current, yielding a highly stable baseline response with a relative standard deviation (RSD) of 2.33%. The corresponding dose-response behavior (Figure 6.iv.) generated the linear regression:

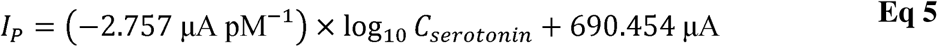

exhibiting a weak correlation coefficient (R^2^ = 0.20111). The current response profile as a function of target concentration is further summarized in the bar graph presented in Figure 6.iii. Based on these measurements, the normalized sensitivity of the PDP-NIP/SPCE was determined to be 21.88 μA (log pM)□¹ cm□². This drastically attenuated sensitivity and the absence of profound signal suppression definitively confirm that the PDP-NIP matrix lacks the structurally complementary cavities required for target sequestration. Consequently, analyte interaction is restricted to weak, non-specific superficial adsorption, firmly establishing the essential role of the imprinting process in the high-affinity performance of the PDP-MIP/SPCE.

**Fig. 6.**
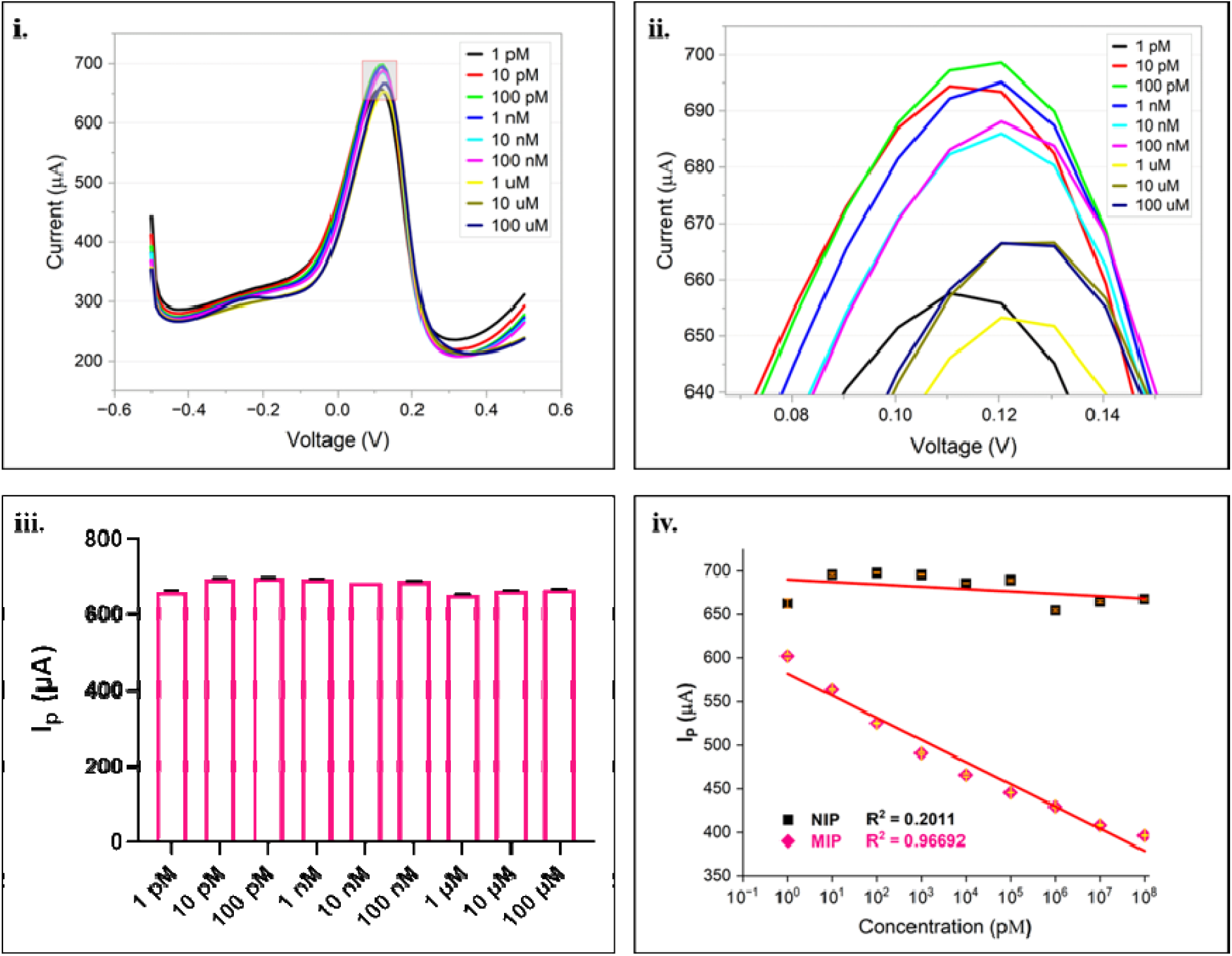
Electrochemical response and control analysis of the non-imprinted polymer (NIP)/SPCE sensor toward Serotonin. (i) Differential pulse voltammetry (DPV) responses of the NIP-modified electrode toward increasing concentrations of Serotonin (1 pM–100 µM), showing limited variation in peak current. (ii) Enlarged view of the DPV responses highlighting the minimal concentration-dependent change in signal. (iii) Corresponding bar plot of peak current responses across concentrations, indicating weak analytical sensitivity and poor discrimination. (iv) Calibration plot comparing MIP and NIP responses, where the NIP exhibits negligible correlation (R² = 0.2011) compared to another MIP (R² = 0.9669), confirming the absence of specific binding sites and validating the role of molecular imprinting in selective serotonin recognition.

#### 3.3.5 Precision and Stability Assessment of the Sensor

To assess the robustness of the engineered sensing platform, its repeatability, reproducibility, and long-term stability were systematically evaluated (Figure 7). Operational repeatability was assessed by performing nine consecutive DPV measurements on the same electrode at a fixed serotonin concentration of 100 nM. Following the expected initial current attenuation upon analyte binding, the sensor maintained a highly stable signal profile across the subsequent eight scans, yielding an RSD of 0.45% (n = 9). To evaluate the consistency of the fabrication process, the reproducibility was assessed across five independently prepared electrodes (E1–E5). The resulting DPV responses demonstrated consistent peak currents with a low RSD of 6.3%, confirming the batch-to-batch reproducibility of the electrode modification procedure.

**Fig. 7.**
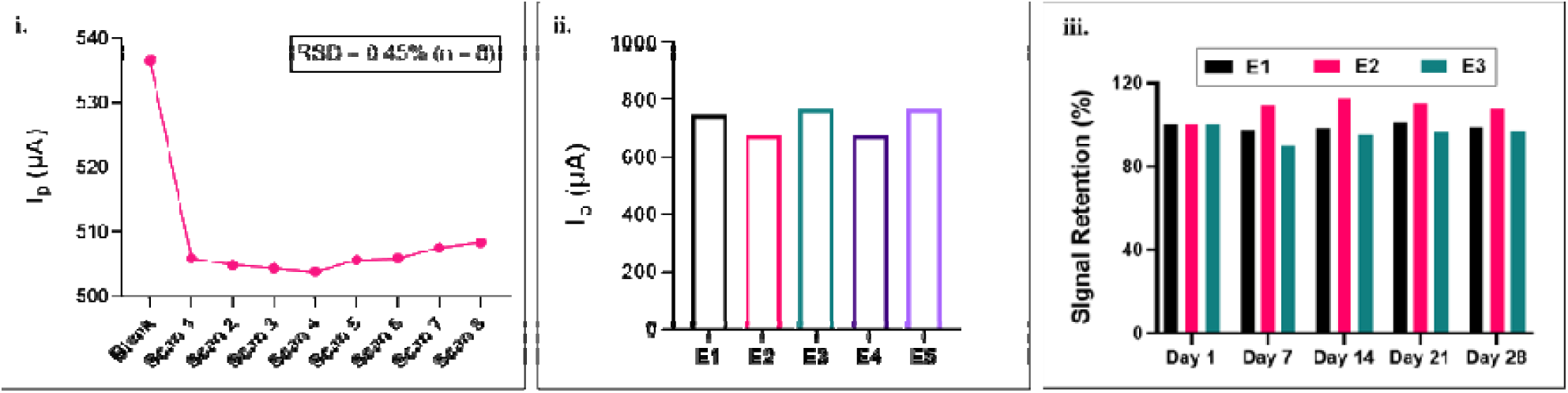
Repeatability, reproducibility, and storage stability of the PDP-MIP/SPCE sensor. *(i)* Repeatability study of the sensor response toward Serotonin measured over successive runs using the same electrode, demonstrating consistent peak current with minimal variation. *(ii)* Reproducibility analysis across independently fabricated electrodes, showing comparable responses and confirming fabrication consistency. *(iii)* Shelf-life evaluation of the sensor over 28 d ys, expressed as signal retention (%), indicating stable performance with negligible degradation under storage conditions. Error bars represent standard deviation (n = 3).

Furthermore, the practical shelf-life of the platform was monitored by tracking the signal retention of three separate sensors (E1, E2, and E3) over 4 weeks (Figure 7.iii.). Measurements taken at intervals of 1, 7, 14, 21, and 28 days revealed excellent storage stability, with the sensors exhibiting minimal deviation and successfully retaining between roughly 90% and 110% of their initial signal responses. Collectively, these remarkably tight variances validate the structural robustness and long-term operational viability of the MIP platform for consistent analytical monitoring.

#### 3.3.6 Interferent Study

Given that non-specific interactions critically compromise analytical accuracy, rigorous selectivity is a fundamental prerequisite for robust electrochemical sensing platforms. To critically evaluate the anti-interference capabilities of the engineered PDP-MIP/SPCE, specificity assays were conducted using DPV in PBS containing [Fe(CN)_6_]^3−/4−^. The sensing interface was challenged with a panel of relevant physiological interferents, specifically glucose, glycine, tryptophan, urea, ascorbic acid, and histamine. Each competing species was introduced at an elevated concentration of 1mg/mL with 100 nM Serotonin to evaluate the sensor’s performance beyond the dynamic analytical range of the target. As shown in Figure 8.i, the targeted addition of Serotonin resulted in a significant attenuation of the voltametric cathodic peak current. Crucially, when co-interferents were introduced alongside Serotonin, they yielded no statistically significant (ns) deviations from the baseline target signal. This high selectivity in complex mixtures is quantified in Figure 8.ii, where the sensor’s signal retention remained exceptionally stable, tightly confined between ∼99% and ∼101%. Control experiments using the **non-imprinted polymer (NIP)** (Figure 8.iii) showed a static, non-discriminatory response across all analytes, notably lacking the signal drop characteristic of Serotonin. The NIP electrode maintained a consistent profile toward various interfering species with an overall **RSD of 2.63%**, confirming uniform non-specific interactions. These results validate that the sensor’s high selectivity stems exclusively from the engineered **MIP cavities** tailored for Serotonin.

**Fig. 8.**
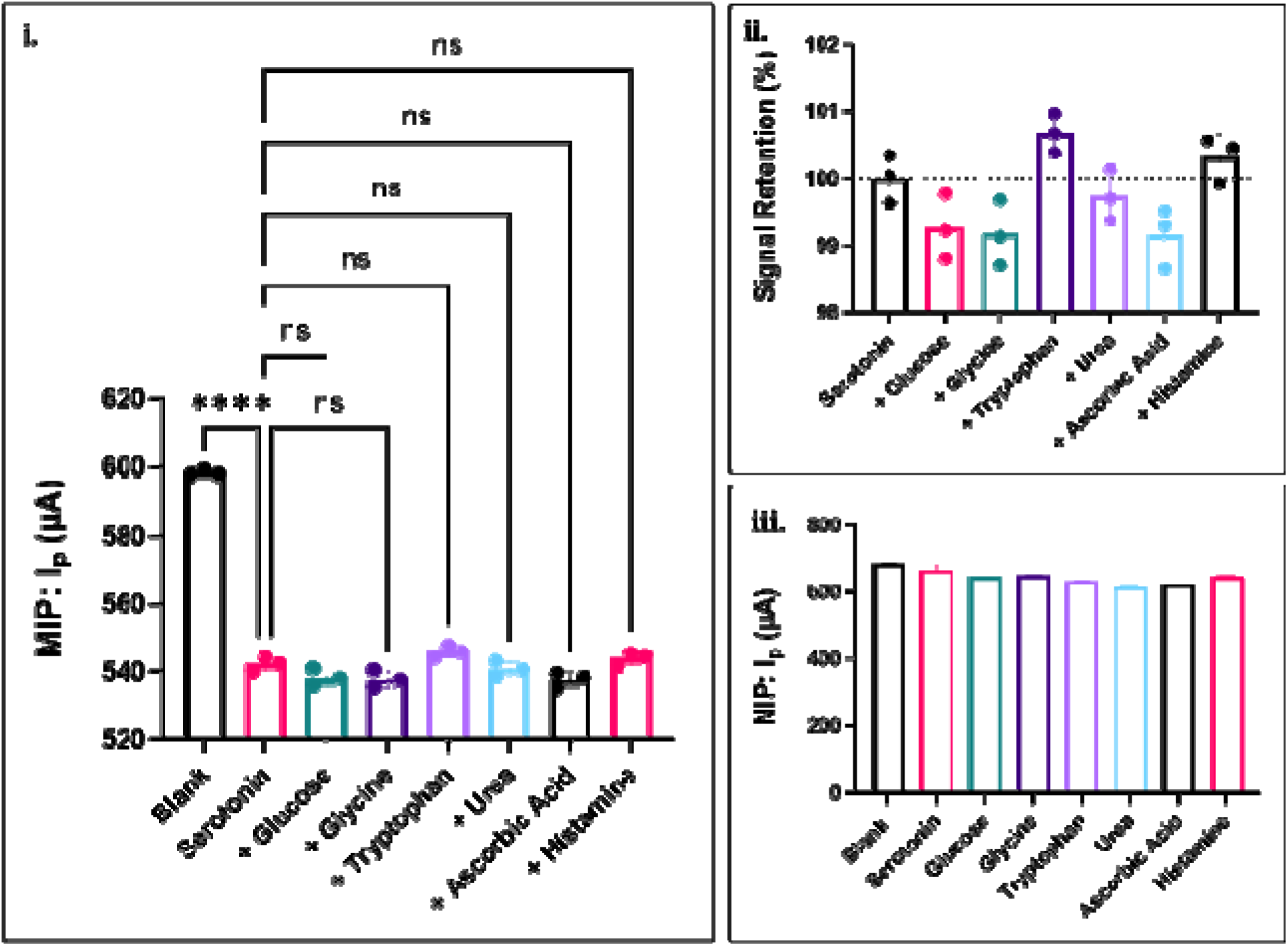
Selectivity and interference analysis of the PDP-MIP/SPCE sensor in comparison with the non-imprinted control. *(i)* DPV peak current responses of the MIP-modified electrode toward Serotonin in the presence of co-existing interfering species, demonstrating preferential recognition of the target analyte. *(ii)* Signal retention (%) of the MIP sensor in the presence of interferents, indicating minimal interference and high selectivity under complex conditions. *(iii)* Interference study performed using the NIP-modified electrode, showing comparatively reduced selectivity and higher susceptibility to non-specific interactions. Error bars represent standard deviation (n = 3).

#### 3.3.7 Real Sample Analysis

The performance of the PDP-MIP/SPCE in a biologically relevant matrix was evaluated using artificial human serum spiked with known serotonin concentrations, spanning 1 pM to 100 μM. Measurements were conducted on both undiluted serum and a 1:10 diluted preparation to assess the impact of matrix complexity on sensor response. Consistent with the established signal transduction mechanism, escalating serotonin concentrations produced a proportional attenuation of the DPV Faradaic peak current in both serum conditions (Fig. 9).

**Fig. 9.**
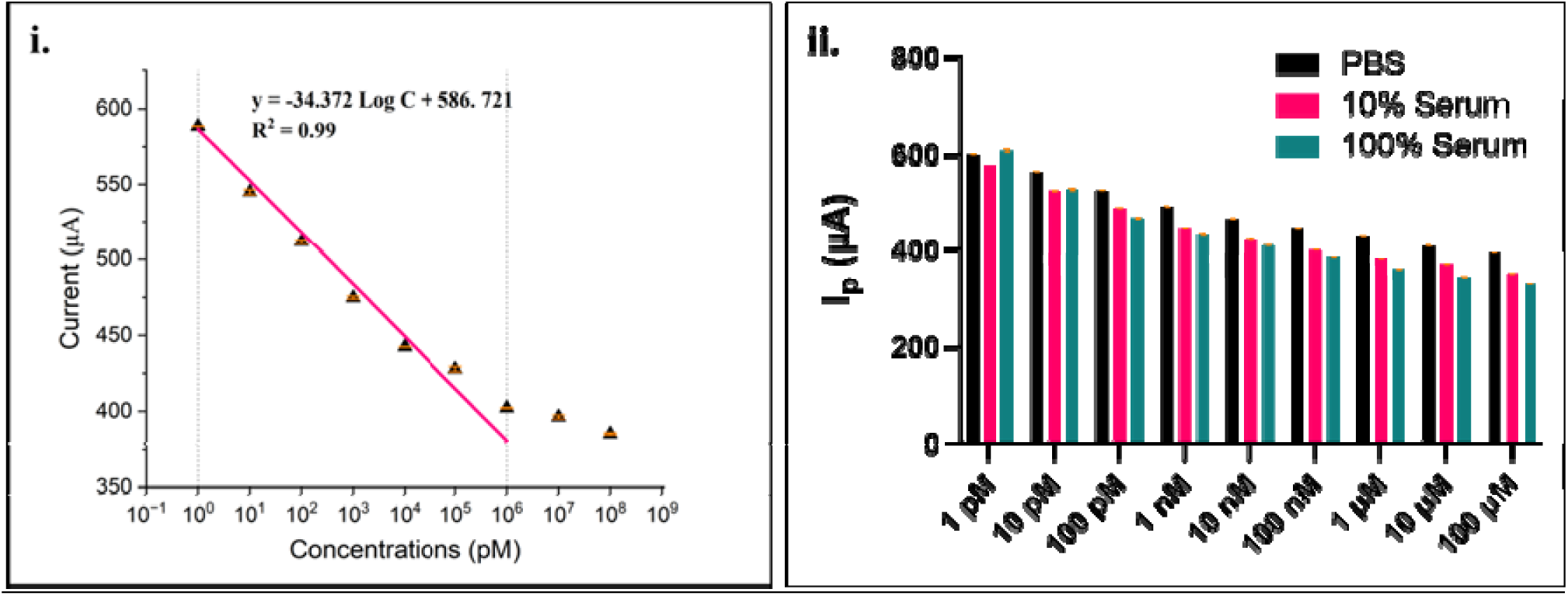
Electrochemical detection of Serotonin in spiked serum samples and comparison with buffer responses. *(i)* Calibration plot of differential pulse voltammetry (DPV) peak current versus logarithmic concentration of Serotonin in spiked serum samples, showing a linear response within the defined concentration range (1 pM–1 µM) (R² = 0.99). *(ii)* Comparative bar plot of DPV peak currents obtained in buffer (pink) and spiked serum samples (teal) across increasing serotonin concentrations, illustrating consistent trends with a slight decrease in signal in serum due to matrix effects. Error bars represent standard deviation (n = 3).

As summarized in Table 2, recovery rates for the 1:10 diluted serotonin-spiked artificial serum ranged from 88.66% to 96.02%, with RSD values between 2.89% and 8.43%. For undiluted samples, recovery ranged from 83.71% to 101.15%, with RSDs between 0.79% and 12.47%. The serotonin LOD in the serum matrix was determined to be 0.12 pM, with a linear dynamic range of 1 pM to 1 μM (R² = 0.99).

**Table 2.** Quantitative determination of Serotonin in spiked human serum samples using the PDP-MIP/SPCE sensor. Analytical performance of the PDP-MIP/SPCE sensor for serotonin detection in spiked human serum samples. Known concentrations of Serotonin were added to diluted serum, and the corresponding differential pulse voltammetry (DPV) peak currents were recorded. The measured responses were used to determine recovery (%) and relative standard deviation (RSD, n = 3), demonstrating the accuracy, precision, and reliability of the sensor in complex biological matrices across a wide concentration range.

| Concentration of Serotonin spiked in the serum | Mean DPV peak current for Serotonin (μA) | Mean DPV peak current of spike-sample (μA) | Relative standard deviation (RSD %) | Recovery (%) |
| --- | --- | --- | --- | --- |
| **Serum Dilution: 1:10** |  |  |  |  |
| 1 pM | 601.83 | 577.90 | 2.89 | 96.02 |
| 10 pM | 564.13 | 523.87 | 5.22 | 92.86 |
| 100 pM | 525.47 | 487.97 | 5.17 | 92.86 |
| 1 nM | 490.90 | 447.13 | 6.61 | 91.08 |
| 10 nM | 465.90 | 423.43 | 6.77 | 90.89 |
| 100 nM | 445.50 | 400.37 | 7.62 | 89.87 |
| 1 μM | 429.63 | 382.57 | 8.26 | 89.04 |
| 10 μM | 408.30 | 370.73 | 6.77 | 90.80 |
| 100 μM | 396.37 | 351.43 | 8.43 | 88.66 |
| **Serum Dilution: Undiluted** |  |  |  |  |
| 1 pM | 601.83 | 608.76 | 101.15 | 0.79 |
| 10 pM | 564.13 | 526.4 | 93.31 | 4.88 |
| 100 pM | 525.47 | 466.46 | 88.77 | 8.36 |
| 1 nM | 490.90 | 434 | 88.41 | 8.72 |
| 10 nM | 465.90 | 409.36 | 87.87 | 9.15 |
| 100 nM | 445.50 | 385.5 | 86.53 | 10.3 |
| 1 μM | 429.63 | 359.3 | 83.63 | 12.67 |
| 10 μM | 408.30 | 343.53 | 84.14 | 12.14 |
| 100 μM | 396.37 | 331.8 | 83.71 | 12.47 |

## 4. Cross-Platform Validation and Point-of-Care Applicability

Cross-platform validation was conducted by performing DPV-based serotonin measurements using two potentiostat systems in parallel: the PalmSens EmStat4S, a benchtop-class portable potentiostat, and the PalmSens Sensit Smart, an ultra-compact module that connects directly to the USB-C port of a standard smartphone and is controlled via a dedicated mobile application PSTouch (Fig. 10). All measurements were performed on the same batch of PDP-MIP electrodes under identical electrolyte conditions and matched DPV parameters. The calibration curves obtained on both platforms exhibited nearly overlapping response profiles across the tested concentration range. No significant shift in peak potential was observed between the two systems. Cross-platform correlation analysis yielded R² = 0.9967 and a slope of 1.023, confirming near-unity proportionality between the two instrumental outputs. These results confirm that sensor performance is governed by the PDP-MIP interface rather than the instrumentation, establishing that the Sensit Smart–smartphone configuration can reliably replicate laboratory-grade measurements. Collectively, this demonstrates the seamless translation of the proposed sensor from benchtop validation to portable, point-of-care deployment.

**Fig. 10.**
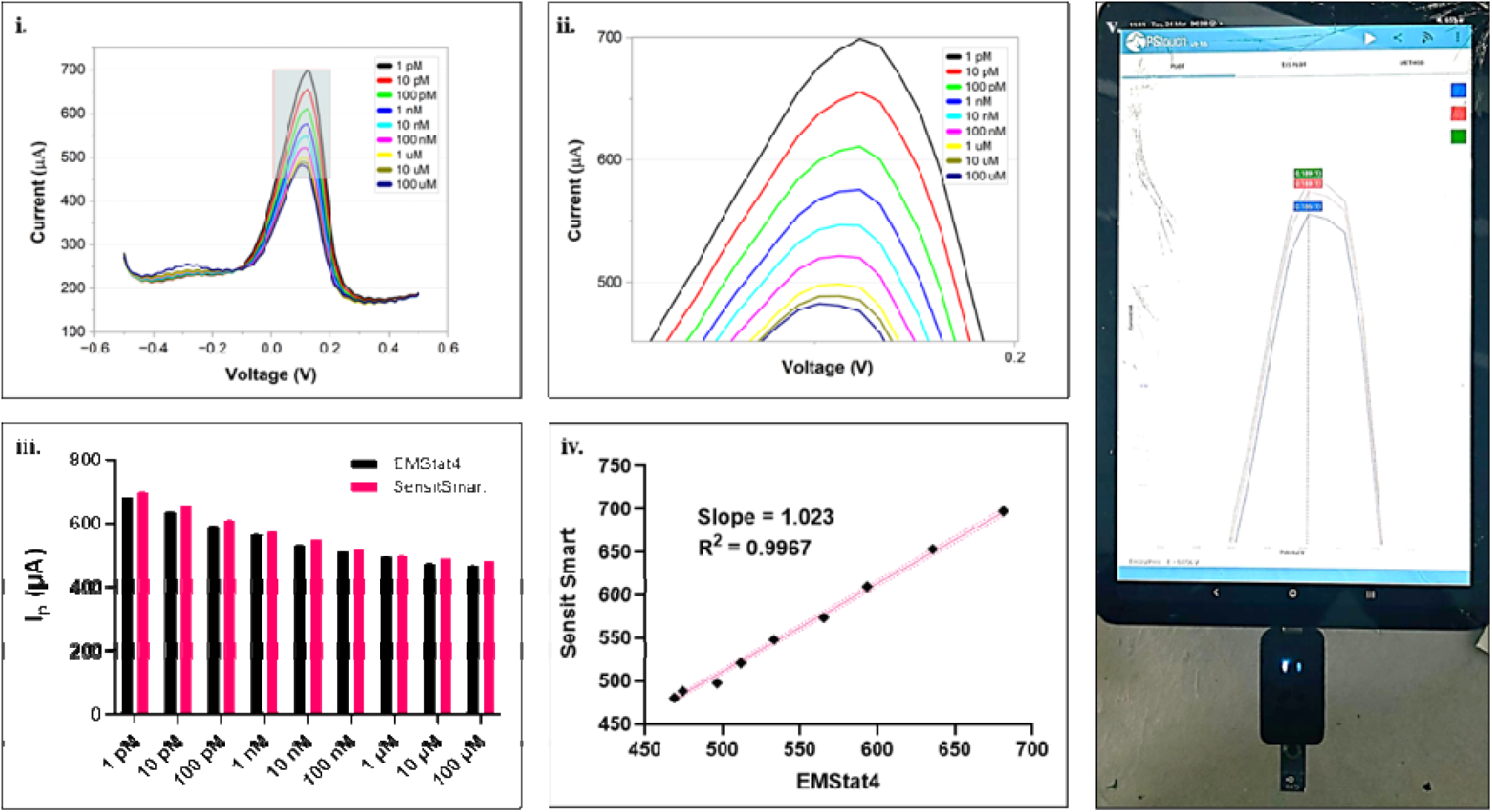
DPV-based Serotonin sensing using a portable platform and validation across independent electrochemical systems. (i) Differential pulse voltammetry (DPV) responses of the MIP-modified electrode toward increasing concentrations of Serotonin (1 pM to 100 µM) recorded using the Sensit Smart platform, showing a concentration-dependent modulation in peak current. (ii) Magnified view of the DPV responses highlighting the systematic variation in peak intensity with analyte concentration. (iii) Corresponding peak current responses plotted as a function of serotonin concentration, demonstrating the quantitative sensing behavior and dynamic range. (iv) Cross-platform validation of the sensor performance using EmStat4 and Sensit Smart devices, exhibiting a strong linear correlation (R² = 0.9967) and slope of 1.023, confirming excellent agreement between platforms. (v) Photograph of the portable Sensit Smart setup used for on-site electrochemical measurements.

## 5. Conclusion

In summary, this work demonstrates the successful development of a portable molecularly imprinted electrochemical sensing platform for the selective and sensitive detection of serotonin using a polydopamine-based MIP layer integrated directly onto screen-printed carbon electrodes. The electropolymerized MIP architecture served as an artificial recognition interface by generating serotonin-complementary rebinding cavities after template removal, thereby enabling selective molecular recognition in both buffered and biologically relevant media. Under optimized conditions, the sensor exhibited a broad linear dynamic range from 10 pM to 10 μM in phosphate buffer, with a correlation coefficient of R² = 0.974, an ultralow limit of detection of 0.16 pM, and a normalized sensitivity of 212.9 μA (log pM)□¹ cm□². Electrochemical kinetic evaluation further confirmed a surface-confined and adsorption-controlled sensing process with rapid electron-transfer characteristics, supporting the efficient interaction between serotonin and the imprinted electrode interface. The analytical reliability of the developed MIP/SPCE platform was confirmed through comprehensive selectivity, reproducibility, repeatability, stability, and real-matrix validation studies. The sensor demonstrated excellent discrimination against physiologically relevant interferents, maintaining signal retention between 99% and 101%, which confirms the high chemical selectivity imparted by the imprinted recognition sites. In addition, the platform showed excellent operational repeatability with an RSD of 0.45%, satisfactory inter-electrode reproducibility with an RSD of 6.3%, and robust 28-day storage stability with 90–110% signal retention. Importantly, validation in spiked artificial serum yielded an even lower LOD of 0.12 pM, along with recovery values of 88.66–96.02% across the tested concentration range, demonstrating that the sensor retains quantitative performance in a protein-rich biological matrix without the need for extensive sample pretreatment. To further establish its suitability for decentralized analysis, the sensing platform was validated using a smartphone-operated miniaturized potentiostat and compared with a laboratory-grade portable potentiostat. The strong cross-platform correlation of R² = 0.9967 and slope of 1.023 confirmed analytical equivalence between the two systems, indicating that the engineered MIP/SPCE interface governs sensor performance independently of instrumentation. Collectively, the disposable SPCE strip, miniaturized potentiostat module, and smartphone-based readout constitute a practical point-of-care analytical system for on-site serotonin monitoring. Overall, the proposed platform offers a simple, cost-effective, robust, and field-deployable strategy for decentralized serotonin detection in clinically relevant samples and may be extended toward the development of MIP-based sensing devices for other neurochemical and disease-associated biomarkers.

## Notes

The authors declare that they have no known competing financial interests or personal relationships that could have appeared to influence the work reported in this paper.

## Acknowledgments

A.K.Y. acknowledges the financial support received from the Anusandhan National Research Foundation (ANRF) under the ANRF-National Postdoctoral Fellowship (PDF/2025/007589). D.B. thanks SERB, GoI, for the Ramanujan Fellowship, DST-Nidhi Prayas for the start-up grant, and Gujcost-DST, GSBTM, BRNS-BARC, and HEFA-GoI, MoES, for STARS for research grants.

## CRediT authorship contribution statement

**Harshit Borasi:** Conceptualization, Methodology, Investigation, Formal Analysis, Data Curation, Writing – Original Draft. **Bhagyesh Parmar:** Methodology, Investigation, Writing – Review and Editing. **Pranjal Aggrawal:** Methodology. **Dhiraj Bhatia:** Funding acquisition, Project administration, Resources, Supervision, Validation, Writing – review & editing. All authors have read and approved the final version of the manuscript. **Amit K. Yadav:** Conceptualization, Methodology, Supervision, Writing – original draft, Project administration, Supervision, Validation, Visualization, Writing – Review and Editing.

## Data Availability Statement

No new data were generated or analyzed in this study.

## Declaration of Generative AI and AI-assisted technologies in the writing process

During the preparation of this work, the author(s) used ChatGPT4 to improve readability and language. After using this tool/service, the author(s) reviewed and edited the content as needed and take(s) full responsibility for the content of the published article.

